# scPlOver: inferring DNA content from amplification-free single-cell WGS using fragment overlaps

**DOI:** 10.64898/2026.05.06.722337

**Authors:** Matthew A. Myers, Gryte Satas, Data Science TeamLab, Sohrab P. Shah, Andrew McPherson

## Abstract

Correctly inferring copy-number aberrations from single-cell DNA sequencing data requires estimating cellular DNA content, which is unidentifiable from read counts alone. In tagmentation-based sequencing, each fragment represents a distinct DNA molecule, thus fragment overlaps provide an orthogonal signal for copy number. We present a theoretical model of fragment overlaps as a function of copy number and coverage and introduce scPlOver, a method that uses this model to infer DNA content. scPlOver outperforms existing approaches on simulated and experimental datasets and identifies thousands of ovarian cancer cells with higher DNA content than previously estimated across a cohort of 41 patients.

## 1 Background

Cancer is an evolutionary process in which cells accumulate mutations over time. One of the most frequent types of mutations is somatic copy-number aberrations (CNAs), changes in the number of copies in a particular genomic region [1]. Individual CNAs can serve as driver events that underlie or improve tumor cell fitness [2], and even the chromosomal instability that causes the accumulation of CNAs can itself serve as a driver of cancer progression [3]. Inherited through cell divisions, CNAs underlie tumor heterogeneity and can be used as indicators of evolution in phylogenetic inference [4–6]. A related type of mutation that is prevalent in some cancer types is *whole-genome doubling* (WGD), in which a tumor cell doubles the number of copies of every chromosome simultaneously (typically via a failure during cell division [7]). While estimates from bulk data [1] suggest that WGD affects ≈37% of primary tumors and a larger fraction of metastases (ranging from 2% to 96% by cancer type [8]), recent studies using single-cell whole-genome sequencing (scWGS) showed evidence that in some tumors, ongoing WGD affects all patients, at least in a small number of cells [9]. Indeed, WGD itself increases the rate of CNA in tumors [9, 10], which may drive further heterogeneity. Thus, assessing the CNAs and WGD status of individual cells is crucial for understanding tumor evolution.

While many methods infer CNAs from both bulk [11–18] and single-cell [19–22] DNA sequencing data, they are unable to resolve tumor ploidy directly. These methods typically leverage the assumption that read counts are directly proportional to integer copy numbers, and seek to assign copy numbers to bins in order to best fit this proportionality (among other objectives, such as minimizing the number of distinct segments, and/or fitting allelic signals such as B-allele frequency [BAF] to infer allele- or haplotype-specific copy numbers). However, due to this proportionality, read counts and allele ratios^1^ alone are insufficient to distinguish between a copy-number profile and an alternative profile in which all states are multiplied by some positive integer *c* (Fig. 1A). As a result, copy-number inference methods typically return the solution with lower average copy number (i.e., *c* = 1), which usually corresponds to a more parsimonious and biologically plausible scenario with fewer ancestral CNAs. In practice, *c* = 2 could indicate a cell in G2 phase of the cell cycle, or a cell that recently underwent whole-genome doubling. Thus, these methods would effectively miss a recent WGD event resulting in a perfect doubling of the chromosomes.

**Fig. 1:**
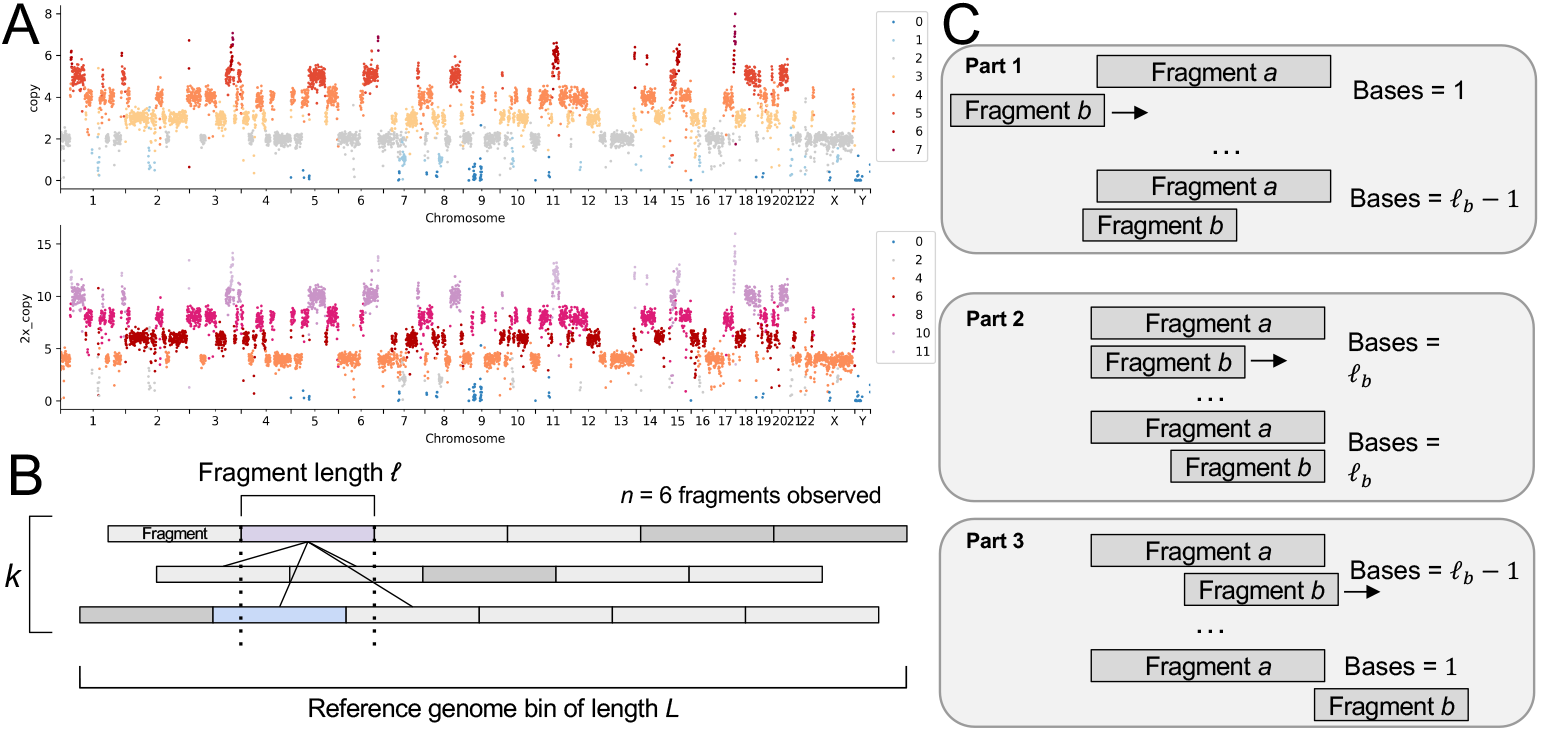
Addressing DNA content non-identifiability using fragment overlaps. **A**. Example copy-number profiles corresponding to the simplest sufficient (*c* = 1) fit of integers to the observed read counts (top) and a doubled profile (*c* = 2, bottom) which could be a G2 cell or cell that recently underwent WGD. **B**. Illustration of fragment overlap model and notation. Rectangles depict tiling of genome copies in a specific bin of length *L* into possible fragments of (uniform) length *𝓁*. Solid lines connect purple-shaded “index” fragment to other tiles which would represent overlaps if they were observed. Shaded fragments indicate randomly sampled observations. **C**. Derivation of the expected number of overlap bases per fragment overlap.

Two recent methods for inferring copy number from scWGS data [23, 24] have emerged that resolved this ambiguity by incorporating fragment overlaps as an orthogonal signal. In certain scWGS technologies such as DLP+ [25] and ACT [26], individual fragments of DNA are tagmented before amplification – so each represents a unique segment of a DNA molecule. The two ends of each fragment are then sequenced via paired-end DNA sequencing, and the genomic location of the fragment can be recovered as the sequence of the reference genome interposed between the two properly paired reads. When two such fragments *overlap* (i.e., align to the same reference positions), this indicates that they must have been drawn from distinct genomic copies of the region. As a result, the number of overlapping fragments is related to copy number: in regions with more copies, there are more fragment pairs that align to the same positions in the reference genome. In Schneider et al. (method scAbsolute) [23], the authors derive statistics for each cell that they show to be related to DNA content (the average copy number of a cell after accounting for cell cycle) using data with experimentally controlled DNA content. Similarly, Kolbeinsdottir et al. (method ASCENT) [24] propose a statistic that uses overlaps to indicate whether an inferred copy-number profile should be doubled or halved. However, both approaches stop short of a comprehensive model for fragment overlaps that explains the expected relationship between fragment overlaps and DNA content while accounting for covariates such as sequencing coverage and fragment length, which obscures their assumptions and limits their interpretation and generalization.

In this work, we present a model of fragment overlaps and derive the expected number of fragment overlaps as a function of DNA content, number of fragments, fragment length, and bin size. We extend this model to the number of total overlap bases (i.e., the total number of reference bases shared by a pair of overlapping fragments), and we show on simulated data that these predictions are accurate and their accuracy improves with sequencing coverage. Building on this model, we present a method scPlOver for inferring DNA content (the average number of copies of the genome in a cell, including cell-cycle effects) from amplification-free DNA sequencing data. scPlOver identifies the DNA content that best fits the data using a Gaussian HMM with emission means closely tied to the expected number of sequencing reads and fragment overlap bases expected by the model for each copy-number state. We apply scPlOver together with the ploidy adjustment modules from ASCENT and scAbsolute to previously published datasets with experimentally sorted G1 and G2 cells, and show that scPlOver outperforms the other approaches and accurately distinguishes cell cycle states with accuracy increasing as a function of sequencing coverage. In previously published high-grade serous ovarian cancer (HGSOC) data, we analyze 24 683 high-quality tumor cells from 41 patients and identify 6 324 cells with at least double the DNA content implied by their originally inferred ploidy.

## 2 Results

### 2.1 scPlOver

We developed a theoretical model for fragment overlaps that enables principled estimation of DNA content from tagmentation-based sequencing data, and derived closed-form equations for the expected number of overlaps and overlap bases (Section 5.1, Figure 1). Consider a genomic bin of length *L* at copy number *k*. Under tagmentation-based sequencing, each copy of the bin is fragmented into an average of *L/𝓁* tiles for a total of *k*(*L/𝓁*) fragments across all copies. When *n* tiles (fragments) are sampled via sequencing, two fragments may overlap if they derive from distinct copies of the same region. For large *L* and *n*, the expected number *X* of such overlaps is

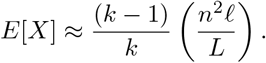

The exact equation is derived in the Methods (Section 5.1). We further extend this model to estimate overlap bases (i.e., the number of base positions in the reference genome shared by each pair of overlapping fragments), which provide a higher-resolution signal than overlap counts alone.

To apply this model to data, we developed a method scPlOver (<u>s</u>ingle-<u>c</u>ell <u>Pl</u>oidy from read <u>Over</u>laps), which infers per-bin copy number and overall DNA content for each cell using a Gaussian HMM whose emission means are derived from the model’s predictions (Section 5.2). We use the term *DNA content* to refer to the average number *d* of copies of the genome present in a cell at time of sequencing, regardless of its cell cycle state. This is distinct from *ploidy*, which refers specifically to copy number in G1 phase and is the quantity most methods estimate.^2^ scPlOver takes as input read counts and overlap bases for each bin, as well as an initial copy-number profile (the assigned states are used in bin filtering and GC correction, and the set of permitted hidden states is constrained to be a function of the original set of states in the profile). We fit scPlOver to each cell using the Baum-Welch algorithm, and we use the Viterbi algorithm to identify the most likely copy-number profile.

### 2.2 Model predicts fragment overlaps and overlap bases from simulated data

To evaluate how well our model predicts fragment overlaps in error-free data, we simulated fragment overlaps at the bin level by tiling each of *k* copies into fixed-length fragments and sampling these fragments uniformly at random (see Section 5.6 for details). Note that this generative process does not explicitly encode our model of the expected number of overlapping fragments. We simulated 8 155 bins with copy number *k* ∈ {1, …, 8} copies, bin length *L* = 500 000, total fragments *F* ∈ [500, 10 000] total fragments, and fragment length *𝓁* ∈ [50, 150]. We found that the predictions from our model were highly accurate in terms of both overlaps (average error 0.79%) and overlap bases (average error 0.68%), and became more accurate as the proportion of fragments sampled increased, analogous to increased sequencing coverage (Fig. 2A-B). For reference, a typical 500-kilobase bin in DLP+ sequencing data may have 500-1000 fragments of average length 150, which would represent 7.5%-15% of possible fragments if the region were diploid (*k* = 2). We repeated these tests on simulated data with variable fragment lengths drawn from an empirical distribution (Section 5.6.2), simulating 6 481 bins each with copy number *k* ∈ {2, …, 8}, total fragments *F* ∈ [200, 10000] sampled from a distribution with mean fragment length 319, and 20 random seeds per parameter setting. Instances where the total length of sampled fragments would have exceeded the available genome length were excluded, so fewer instances with large *F* relative to *k* were included. Results with variable fragment lengths were largely the same as those with fixed lengths (Fig. 2C-D): average error was *<*1% for both overlaps and bases across all simulated instances.

**Fig. 2:**
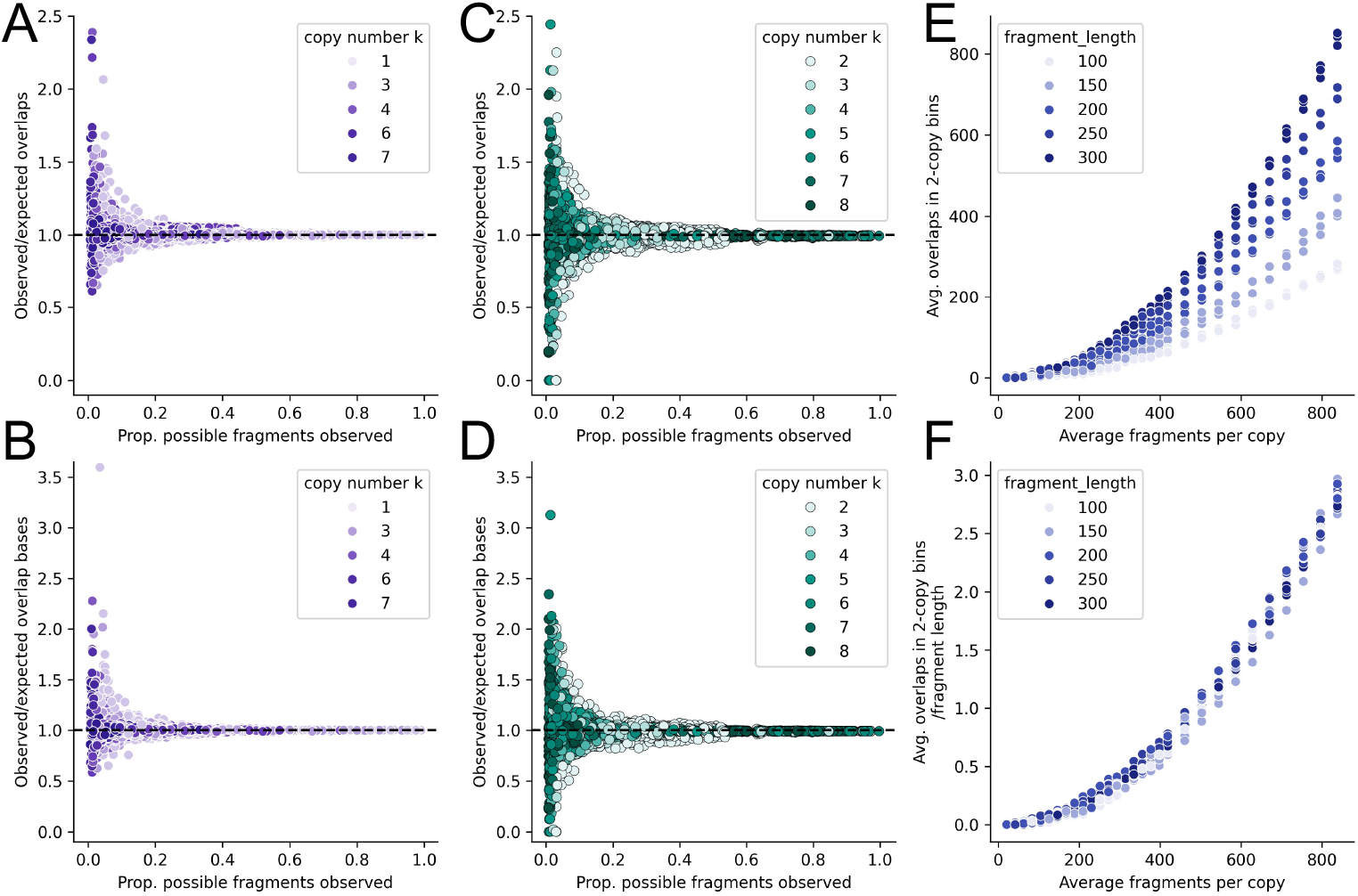
Testing model of fragment overlaps using simulations. **A-B** use fixed-length fragments, **C-D** use empirical fragment lengths. **A**. Ratio of observed to expected fragment overlaps (y-axis) vs. proportion of all possible fragments observed (x-axis) in simulated fragments from individual bins, colored by number *k* of copies. **B**. Ratio of observed to expected overlap bases in the same simulated bins as A. **C**. Same as A except with varying fragment lengths sampled from an empirical distribution. **D**. Same as B except with varying fragment lengths sampled from an empirical distribution. **E**. Average number of overlaps in copy-2 bins (y-axis) in simulated genomes as a function of the average number of fragments per copy (x-axis). **F**. Average number of overlaps in copy-2 bins divided by average fragment length (y-axis) in simulated genomes as a function of the average number of fragments per copy (x-axis).

We also examined the scAbsolute model in the context of simulated data. In Schneider et al. [23], the authors derive two statistics to represent each cell: the average number of reads (fragments) per copy as an analog for sequencing coverage, and the “read density” which is described as the average fraction of overlapping reads (fragments) in regions predicted to have copy number 2 (see Supplement section S3.1 for details). The authors propose regressing read density against fragments per copy in cells of known DNA content, and comparing new cells to this curve: cells with underestimated DNA content should be significantly above this curve. However, this read density measure fails to account for varying fragment length between cells in the same sample: our model predicts that the number of overlaps is roughly linear in the fragment length, so bins with the same state and number of fragments but longer fragments will have higher overlaps in copy-number state 2 regions (y-axis of proposed curve), despite the same number of fragments per copy (x-axis of proposed curve). As a result, differences in average fragment lengths could be spuriously interpreted as differences in DNA content between cells. This prediction was supported by the simulated data which showed curves that varied widely by fragment length (Fig. 2E). scPlOver offers a simple remedy of dividing the average overlaps in copy-number state 2 bins by fragment length, which largely accounted for fragment length in simulated data and produced a single curve that could be used as a reference (Fig. 2F).

### 2.3 scPlOver accurately infers DNA content from experimentally sorted data

To evaluate the performance of scPlOver, we applied scPlOver and the DNA content prediction modules from ASCENT [24] and scAbsolute [23] to 2 823 cells from three DLP+ scWGS datasets that were each generated under experimental conditions designed to control the DNA content in each cell: cells sorted using the FUCCI system [23] (*n* = 255 G1 cells and *n* = 233 G2 cells, base ploidy approximately 3.15), and a pair of datasets sorted using dual markers (DM) for S-phase and DNA content [27] (tetraploid: *n* = 249 G1 cells with base ploidy approximately 3.75 and *n* = 637 G2 cells; diploid: *n* = 677 G1 cells with base ploidy approximately 2 and *n* = 772 G2 cells) (Fig. 3A-B).

**Fig. 3:**
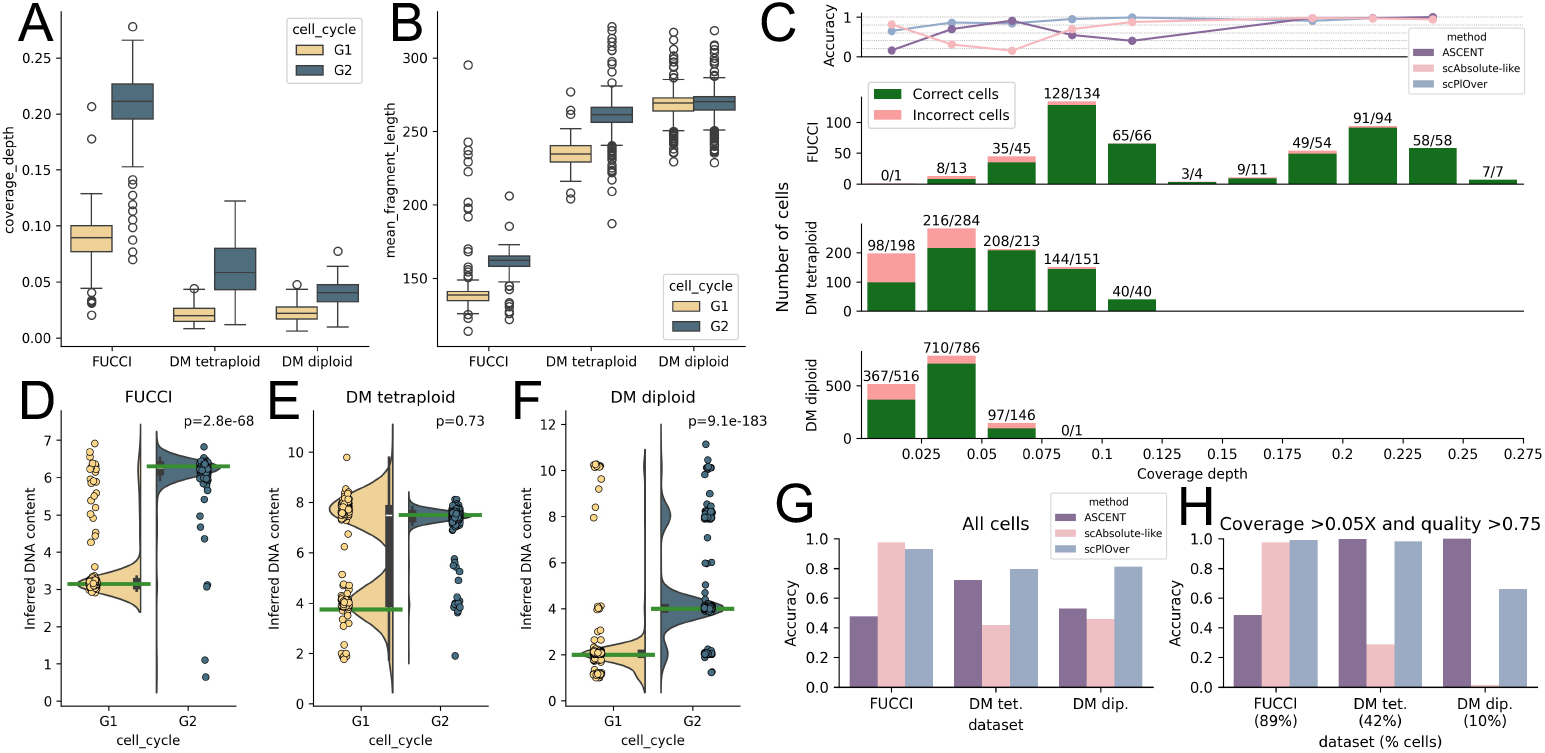
Evaluating scPlOver using data with experimentally controlled DNA content. **A**. Coverage depth (y-axis) and **B**. mean fragment length (y-axis) for each cell in each of the datasets (x-axis) with experimentally controlled DNA content, separated by cell-cycle state. **C**. Accuracy on datasets with experimentally controlled DNA content, partitioned by coverage. Top plot shows accuracy of all methods across all datasets as a function of coverage, including only those bins with at least 15 cells. Below, results of scPlOver are separated by dataset (row) and each bar shows the number of correctly (green) and incorrectly (pink) classified cells and is labeled above with the numbers of correct and total cells, respectively, within the corresponding coverage interval. **D-F**. Inferred DNA content from scPlOver in each experimental dataset, grouped by dataset and condition. Green lines indicate the true DNA content value for the corresponding condition; p-values from two-sided Mann-Whitney U test. **G**. Overall accuracy for each method on each dataset. **H**. Overall accuracy for each method on the subset of each dataset with coverage *>*0.05× and quality *>*0.75. Dataset labels are annotated with the percent of cells in each dataset included in this subset.

We found that despite sequencing coverage as low as 0.006× for some cells, scPlOver classified cells correctly (within 0.5 of the correct DNA content) with an overall accuracy of 82.7%, increasing to 91.2% for cells above 0.05× coverage and further to 95.8% for cells above 0.075× coverage (Fig. 3C). The inferred DNA content from scPlOver typically matched the expected DNA content according to base ploidy and cell cycle phase (Fig. 3D-F). While scPlOver was slightly less accurate on DM tetraploid cells (accuracy 79.7%, mean coverage 0.05×), this accuracy increased to 97.0% when considering only cells with at least 0.05× coverage.

The DNA content prediction module in ASCENT [24] takes per-cell copy-number calls as input and uses an odds ratio to determine whether the inferred ploidy should be kept as-is, halved (DNA content correction factor of 0.5), or doubled (correction factor of 2). ASCENT’s formula assumes that cells have fixed coverage and downsamples input cells to 0.005× to ensure that this is the case; we followed this approach and recomputed overlaps accordingly (see Supplement section S2 for details on the approach used in ASCENT and its application here). Overall, ASCENT achieved an accuracy of 58.3% – while it performed fairly well on the dual-marker datasets (diploid, accuracy 53.0%; tetraploid, accuracy 72.2%), it performed much more poorly on the FUCCI dataset (accuracy 47.7%). ASCENT’s accuracy improved considerably when restricting to the higher-coverage and higher-quality subset of cells: 100% on DM diploid, 99.7% on DM tetraploid, and 48.7% accuracy on FUCCI. Interestingly, although ASCENT achieved a lower accuracy than scPlOver on the DM diploid dataset overall (scPlOver accuracy 81.0%), ASCENT outperformed scPlOver on the high-coverage and high-quality subset of the DM diploid dataset (representing 147*/*1449 = 10% of the dataset).

These copy-number profiles are very simple (Fig. **S4**C), with only ≈6.96% of the autosomal genome altered from diploid. In this setting, the rigidity of the ASCENT approach (which assumes that the correct copy numbers are supplied as input) may represent an advantage.

We were unable to evaluate scAbsolute directly, as the code base does not contain an automatic method for analyzing DNA content using fragment overlaps (see Supplement section S3.2 for details). Instead, we followed the guidelines in the vignette “vignette-ploidy.pdf” in the scAbsolute repository to conduct an analysis that is similar in spirit to the DNA content analysis presented in the original publication. These results are summarized here; see section 5.5 for details on our approach, and Supplement section S3.3 for details on how we arrived at this approach including comparison to published read density values. The manual scAbsolute-like analysis achieved an overall accuracy of 52.3%. While it performed comparably to scPlOver on the FUCCI data which was used in the development of scAbsolute (accuracy 97.5%), it performed much more poorly on the DM datasets (DM tetraploid, accuracy 41.7%; DM diploid, accuracy 45.7%; Fig. 3G). Oddly, its accuracy was lower on the higher-coverage and -quality subset of the DM datasets (DM tetraploid, accuracy 28.9%; DM diploid, accuracy 1.4%; Fig. 3G) than on the full datasets – this is likely due to the fact that the classification on the DM datasets is effectively random (Fig. **S3**E-L). The fragment overlap model in scAbsolute does not take fragment length into account, and the FUCCI dataset has a strong association between fragment length and cell cycle (Fig. 3B); in datasets with weaker associations between fragment length and cell cycle, it does not distinguish between cell cycle states (Fig. **S3**). We elaborate on these results in Supplement section S3.3.

### 2.4 Overlaps reveal cells with elevated DNA content in ovarian cancer cohort

We applied scPlOver to 24 683 high-quality tumor cells from 41 high-grade serous ovarian cancer (HGSOC) patients sequenced using DLP+ scWGS as part of a recent study [9] (mean coverage 0.10×). (We restricted our analysis to this subset of the original 30 260 cells with coverage ≥0.05×, as below this coverage in experimental data scPlOver was less reliable.) In the original analysis of this data, cells were classified using a heuristic based on allele-specific copy number as having 0, 1, or 2 ancestral WGD events (referred to as 0×WGD, 1×WGD, and 2×WGD, respectively). Based on the proportion of cells in the samples classified as having at least 1 WGD (i.e., ≥1 WGD), patients were classified as WGD-low (<15% of cells ≥1×WGD) or WGD-high (50% of cells ≥1×WGD). scPlOver-inferred DNA content largely agreed with the previously inferred ploidy values (73.9% of cells within an absolute difference of 0.5), though agreement varied widely at the patient level, from 0.3% in patient 118 to 100% in patients 026 and 037 (Fig. 4A). The vast majority (98.8%) of disagreements were in the positive direction (i.e., scPlOver inferred a higher DNA content than the original analysis), with only 76 cells inferred to have lower DNA content – and only 7 of those had coverage over 0.1. For cells predicted to have higher DNA content (at least 1.5 times the original ploidy), the increase was typically (91.3%) by a factor of 2 and as such, we refer to these as “doubled” cells.

**Fig. 4:**
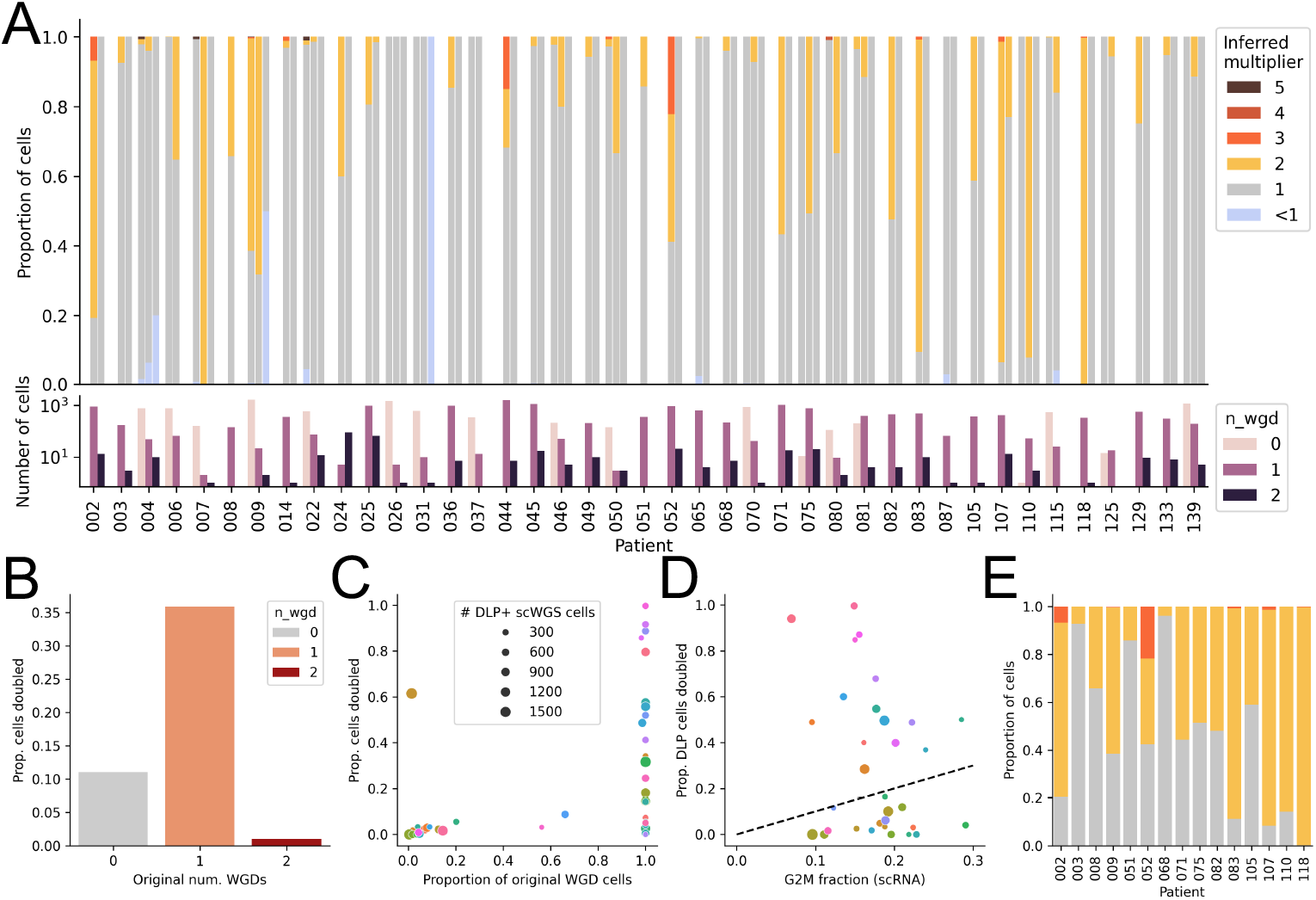
Identifying previously undetected G2 or WGD cells in HGSOC. **A**. Proportion of cells inferred with each DNA content multiplier (colors, above) relative to its original ploidy, separated by patient (x-axis) and grouped by number of ancestral WGD events inferred in the original analysis. Bottom track shows the number of cells with each WGD status in each patient (log-scaled). **B**. Proportion of analyzed cells (y-axis) in each WGD category (x-axis) that were identified as having additional doubling beyond the inferred CN profile. **C**. Proportion of cells identified as doubled (y-axis) and proportion of cells originally called as having at least one ancestral WGD (x-axis) for each patient. Colors indicate patient IDs. Points are scaled by the number of DLP+ scWGS cells for each patient. **D**. Proportion of cells in DLP+ scWGS identified as doubled (y-axis) and proportion of cells in scRNA-seq identified as G2/M phase (x-axis) for the 32/41 patients with matched scRNA-seq data. Colors indicate patient IDs and points are scaled by DLP+ scWGS cell count, same as in C. Dotted line shows *y* = *x*. **E**. For the 15 patients with higher doubling rate than G2/M fraction, proportion of cells with each inferred DNA content multiplier. Bar colors are the same as in A.

To further investigate these doubled cells, we regressed known covariates of genome doubling (mitochondrial DNA (mtDNA) copy number [28] and cell diameter quantified via imaging in the DLP+ spotting nozzle [25]) against the inferred DNA content multiplier (restricting to multipliers with at least 100 cells: 1, 2, and 3) while controlling for patient using ordinary least squares (OLS). We found that both cell diameter and mtDNA copy number had significant positive coefficients (OLS 95% confidence interval lower bound *>* 0), supporting the finding of cells with higher DNA content than originally estimated.

The overwhelming majority of doubled cells were among those originally inferred as 1×WGD: 36% of these cells were inferred to be doubled, as opposed to 11% of 0×WGD cells and 1% of 2×WGD cells (Fig. 4B). At the patient level, only those patients whose cells were nearly all ≥1×WGD in the original analysis had large fractions of doubled cells, with the exception of patient 009 (Fig. 4C). To ascertain whether these doubled cells were likely G2 cells or additional WGD events beyond what was accounted for by the copy number, we examined the cell-cycle phase calls from matched scRNA-seq from the original study for 32/41 patients. We found that these patients were split into two groups: 17/32 patients had proportions of doubled cells lower than their respective G2/M fraction from scRNA. Thus, for these patients the doubling could be attributed to G2 phase cells. This group included all of the WGD-low patients with matching scRNA-seq. However, for the remaining 15 patients, the proportion of doubled cells far exceeded the G2/M fraction in scRNA-seq (Fig. 4D-E).

It is unclear whether these doubled cells represent G2 cells or additional WGD, as while the inferred G2/M fractions in scRNA-seq are low, these cells are known to have abnormal cell cycles [9] and thus may be in an aberrant G2 phase (or G2 arrest) without expressing the typical markers. The 15 patients with high doubling fractions tended to be those patients with high cell-to-cell copy number variability as quantified by significantly higher sample-level normalized rates of arm- and segment-level copy-number gains than the rest of the cohort (*p <* 0.021, two-sided Mann-Whitney U test; Fig. **S5**C-E; see original publication [9] for details on rate computation). This could mean that these cells are unstable and more likely to exhibit G2 arrest, or that they are more likely to undergo additional WGD. Unfortunately, DNA content alone cannot distinguish these two cases.

## 3 Discussion

In this work, we developed scPlOver, a method to infer the DNA content of single cells sequenced using tagmentation-based scWGS. We first derived a theoretical model for fragment overlaps in amplification-free single-cell DNA sequencing data and showed that it accurately predicts overlap counts and overlap bases in simulated data. We then leveraged this model to estimate the DNA content of individual cells. Using three datasets containing experimentally sorted cells with known DNA content, we demonstrated that scPlOver accurately estimates DNA content, with accuracy improving as coverage increases. scPlOver was more accurate on these datasets than other methods that consider fragment overlaps (scAbsolute [23] and ASCENT [24]), though we note that using fragment overlaps to estimate ploidy is a minor component of those methods rather than their primary focus. We next applied scPlOver to the SPECTRUM HGSOC cohort [9], revealing large subpopulations of cells with DNA content exceeding their apparent ploidy by approximately two-fold. Interestingly, these populations were observed in a minority of tumors, characterized by high chromosomal instability and abundant cell-to-cell heterogeneity. We hypothesize that these cells may represent G2 populations substantially larger than expected given matched scRNA-seq measurements, or previously undetected subclones with additional whole-genome doubling.

Certain aspects of the relationship between DNA content and fragment overlaps remain obscure. For example, we currently do not know the cause of the bias factor *δ* by which observed overlap bases exceed expected overlap bases. All sequencing data analyzed in this publication was processed uniformly from FASTQ files (including alignment using `bwa-mem` [29], see Methods for details), so it could be driven by artifacts in alignment. However, this factor was fairly consistent across DLP+ datasets generated by different institutions with vastly different sequencing parameters (fragment length, coverage depth, read length, etc.), suggesting it may be driven by a violation of the uniform sampling assumption caused by tagmentation-related non-uniformity of sequencing coverage. More investigation is needed to identify and address this issue.

Another gap that could be addressed in future methods concerns the bin width *L*. In this work, we use uniform-width 500-kilobase bins and count all fragments within each of these bins, so we use *L* = 500 000 in the model. However, in practice, not all bases in every bin are uniquely mappable, so the effective bin width *L*_*e*_ over which fragments are sampled and overlaps can occur may often be smaller than *L*. Modeling *L*_*e*_ and accounting for variation in coverage breadth across bins (and as a function of the reference genome, sequencing technology, and alignment method and parameters) may enable more accurate inference of DNA content.

## 4 Conclusions

Our findings suggest that fragment overlaps provide a previously underutilized signal for quantifying DNA content in single cells. By enabling more accurate estimation of DNA content, scPlOver has the potential to reveal previously unobserved heterogeneity within cellular populations, improving confidence in the identification of whole-genome doubling, haploidization, and cell-cycle states. These measurements can then be related to other features captured by scWGS, including cell-specific aneuploidies, structural variation, and DNA replication dynamics [27,30], to further understand chromosomal instability and aberrant cell division in tumor populations.

Future work will focus on integrating fragment overlaps into comprehensive models of scWGS copy number inference, enabling joint estimation of DNA content, segmental copy number, and allele-specific signals. Such integration could improve the accuracy and confidence of copy-number calls, enabling more sensitive detection of copy-number variation and associated cellular states. Although our results were demonstrated using DLP+ datasets, scPlOver is in principle applicable to any sequencing technology in which genomic fragments are tagmented prior to amplification and approximately uniform coverage is expected, including ACT [26].

## 5 Methods

In this section, we describe a model for fragment overlaps (and overlap bases) in amplification-free DNA sequencing data that accurately predicts these values in simulated data. Then, we describe our model-informed method scPlOver for inferring DNA content from real data using fragment overlaps. For comparison, in the Supplement we describe the DNA content inference approaches employed in scAbsolute [23] and ASCENT [24] as we understand them (Supplement sections S2 and S3).

### 5.1 Generative model of overlapping fragments

Our goal is to estimate the expected number of fragment overlaps, i.e., the expected number of pairs of fragments that align to at least one of the same reference bases. Consider a region of the reference genome (i.e., a bin) of length *L* that is present in a tumor cell at copy number *k* (after accounting for cell cycle). Each copy of this region in the tumor genome is broken up via tagmentation into DNA fragments with some average length *𝓁*. These fragments represent a tiling of each copy of the tumor genome into an average of *L/𝓁* tiles per copy, for a total of *T* = *kL/𝓁* tiles (Fig. 1B). We observe *n* of these fragments via paired-end sequencing (i.e., we have 2*n* properly paired reads).

We construct a graph (𝒱, ℰ) such that the set 𝒱 of vertices corresponds to the set of tiles, and the set ℰ of edges represents all pairs of overlapping tiles. Let *M* = |ℰ |represent the total number of edges. Assuming each tile overlaps an average of two tiles from every other copy of the genome, and ignoring directionality, 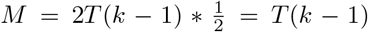 (see Supplement section S1 for derivation). For each edge, *e*, let *I*_*e*_ be an indicator such that *I*_*e*_ = 1 if both adjacent vertices are chosen, and *I*_*e*_ = 0 otherwise. Then, the total number of overlaps *X* =Σ_*e∈ℰ*_ *I*_*e*_. Under the assumption that reads are sampled uniformly at random (which is implicit or explicit in virtually all methods that infer copy number from sequencing data), the expected number of overlaps is as follows.

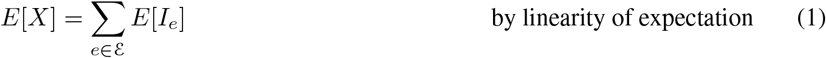

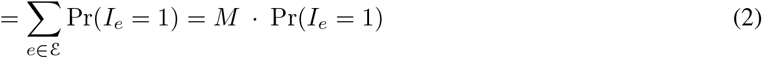

To compute Pr(*I*_*e*_ = 1), we know that the number of combinations of *n* selected tiles is 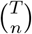. The number of combinations where both vertices from *e* are selected is 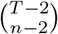. Then,

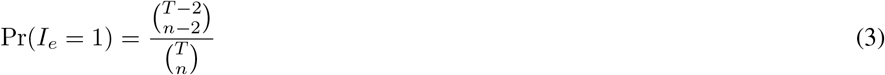

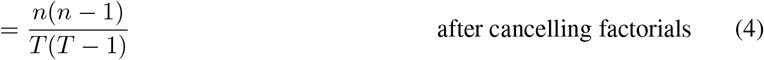

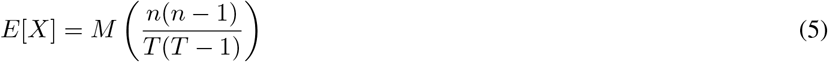

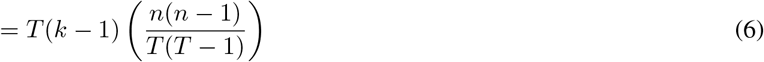

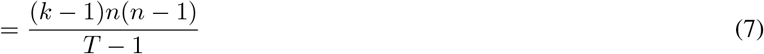

We can extend this model to consider the number of *overlap bases* as well: for each overlap between two fragments, the number of overlap bases is the number of bases in the reference genome that overlap both fragments. To extend the model, we must compute the average number of bases per overlap. We proceed by constructing the piecewise function representing the number of overlap bases at each possible start position (Fig. 1C). Consider two fragments *a* and *b* with lengths *l*_*a*_ and *l*_*b*_ respectively. Fixing the start position of the longer fragment *a* (*l*_*a*_ *> l*_*b*_) WLOG, there are *l*_*a*_ +*l*_*b*_ *−*1 possible start positions for fragment *b* which would result in an overlap between the two fragments. This first section of the function forms an isosceles right triangle with non-hypotenuse side length min(*l*_*a*_, *l*_*b*_) = *l*_*b*_. Starting from the first possible start position with 1 overlapping base, the number of overlap bases increases by 1 with each increase in the start position of fragment *b* until the two fragments are completely overlapping (at which it is equal to min(*l*_*a*_, *l*_*b*_) = *l*_*b*_). The overlap size remains at this value until the end of fragment *a* passes the end of fragment *b* (*l*_*a*_*− l*_*b*_ bases later), then decreases by 1 with each position. The area under this curve is 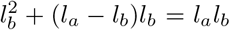, so the average number of bases per overlap is *l*_*a*_*l*_*b*_*/*(*l*_*a*_ + *l*_*b*_ *−*1). In expectation, with an average fragment length of *𝓁*, this quantity is *𝓁*^2^*/*(2*𝓁 −*1). Thus, the expected total number *B* of overlap bases is

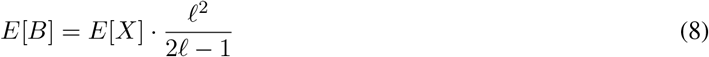

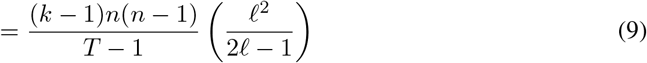

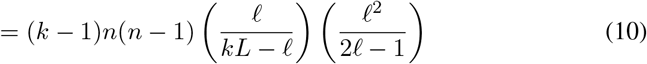

### 5.2 Predicting DNA content using fragment overlaps

Our goal is to identify the most likely DNA content 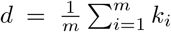(i.e., average copy number accounting for cell cycle) for a particular cell, given read counts *r*_*i*_ and overlap bases^3^ *b*_*i*_ for each bin *i*. We developed scPlOver (<u>s</u>ingle-<u>c</u>ell <u>Pl</u>oidy from read <u>Over</u>laps) which solves this problem by performing a 1-dimensional search over possible values of *d*, evaluating each value by fitting a constrained multivariate Gaussian^4^ HMM to the data. We fix the means *µ*_*r*_(*k*), *µ*_*b*_(*k*) of the Gaussian distribution for each hidden state *k* to the expected number of reads and overlap bases with *k* copies. Let *R* = Σ_*i*_ *r*_*i*_ indicate the total number of sequencing reads from the cell, and let *F* = Σ_*i*_ *n*_*i*_ indicate the total number of distinct fragments from the cell.^5^ Then, let *ρ* = *R*/Σ _*i*_ *k*_*i*_ indicate the average number of reads per copy, and let γ = *F*/Σ _*i*_ *k*_*i*_ indicate the average number of fragments per copy. Leveraging the expected proportionality between reads/fragments and copy number (*r*_*i*_ ≈ *ρk*_*i*_, *b*_*i*_ ≈ γ*k*_*i*_), the expected number of reads *µ*_*r*_(*k*) and overlap bases *µ*_*b*_ (*k*) for a given copy-number state *k* are then:

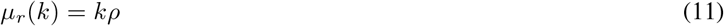

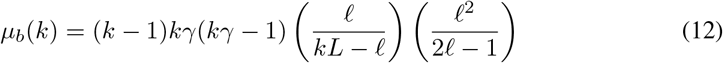

Let *k*_*max*_ indicate the user-specified maximum allowed copy-number state. The transition matrix *T* is initialized with a diagonal structure to penalize changes in copy-number state:

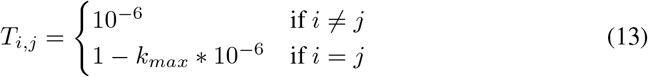

Rather than inferring *k*_*i*_ and *d* jointly, we fix an initial *d* which determines the expected means for each *k*_*i*_, fit the transitions *T* and covariances Σ using the Baum-Welch algorithm, then we marginalize over the copy numbers *k*_*i*_ to compute the likelihood of the data conditioned on *d* using the forward-backward algorithm. We repeat this process for several different values of *d* and return the value of *d* which results in the highest likelihood.

To avoid model selection challenges in which larger values of *d* condense more states to the same magnitude as the data, we restrict the number of permitted hidden states (i.e., *k* values). Specifically, scPlOver takes as input a set *ϕ* of permitted copy-number states and an initial DNA content estimate *d*_0_, and considers only those values of *d* that correspond to integer multiples or divisions of *d*_0_ (with allowed copy-number states corresponding to the same multiplication or division applied to the original copy-number states). For example, given DNA content *d*_0_ and unique states *ϕ* = {0, 1, 2, 3}, we consider also the DNA content 2*d*_0_ with states {0, 2, 4, 6}. In practice, we derive *d*_0_ and *ϕ* from an input copy-number profile.

This approach constrains the solution space while using limited information about the input copy-number profile.

### 5.3 Accommodations for real data

In practice, we modify the Gaussian HMM emission means in three ways to adapt to real sequencing data. First, we observed a bias factor *δ*_*k*_ by which observed overlap bases exceed expected overlap bases in cells with known DNA content, which is fairly consistent across cells, libraries, and datasets. This factor starts at *δ*_2_ = 1.61 and rapidly descends toward *δ*_12_ = 1.08. We derived values for *δ* by analyzing the empirical ratios of observed to expected overlap bases.

The second modification is that we fit constant coefficients *β*_*r*_, *β*_*b*_ ∈ [0.8, 1.2] to permit some flexibility for noise in the data without straying too far from the model-determined expectation. We found this to be useful for lower-coverage data with inconsistent quality, and on experimental data both *β*_*b*_ and *β*_*r*_ approach 1 as coverage increases (Fig. **S5**A-B). In summary, for typical analysis in this publication, the emissions use means 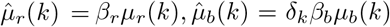 where *δ*_*k*_ is fixed and *β*_*r*_, *β*_*b*_ are learned. For the SPECTRUM dataset with consistent sequencing quality and established cell filtering from the original analysis [9], we fix *β*_*r*_ = 1 and use tighter bounds *β*_*b*_ ∈ [0.95, 1.05]. Whether or not to fit *β*_*b*_ or both *β* parameters, as well as their bounds, are exposed as command-line arguments that can be tuned by the user.

Third, to accommodate the small number of overlap bases observed in regions with *k* ∈ {0, 1} that are not expected by the model, we inflate *µ*_*b*_(0) from 0 to 1, and *µ*_*b*_(1) from 0 to 5.16 (in experimental datasets) or to the empirical mean of state-1 bins (in SPECTRUM datasets).

### 5.4 Preprocessing and parameter settings for analysis

When counting fragment overlaps, we restricted our analysis to fragments with a mapping quality of at least 20 and length at least 50. Using normal cells, we identified a blacklist of 322 bins containing poorly behaving overlap signal (defined as observed overlaps less than 0.5 times or more than 5 times expected) in our data aligned to GRCh37 (see paper repository for list), and removed these bins from consideration. For each cell, we removed bins with no fragments remaining after filtering. We also removed states (and the corresponding bins) with fewer than 50 assigned bins in the input copy-number profile, and any bin outside of 2 \**IQR* from the mean value (in either dimension – reads or overlap bases) for the assigned state. We performed GC correction using modal quantile regression as previously described [9], fitting separate GC curves for reads and overlap bases. Copy-number profiles were inferred using HMMCopy, and only those cells with a quality score^6^ of at least 0.5 were analyzed. To scale modal-regression-corrected values back to the space of the original data, we regressed the corrected values against their expected overlap bases (according to the assigned state in the input copy-number profile, and incorporating *δ*) via first-order least squares regression with no intercept. When scaling overlap bases, to mitigate the effect of high-variance bins, we removed from consideration any bin above the 80th percentile of absolute difference from its expected overlap bases given the state assigned to the bin.

To determine the ground truth DNA content values for experimental datasets, we corrected the modal HMMCopy copy number profile by multiplying its average value by an integer multiple to match the indication for each cell (e.g., the modal HMMCopy profile across the cells in the FUCCI dataset had an average copy number of 3.12, so the G2 cells in the dataset were assumed to have a true DNA content value of 6.24).

### 5.5 scAbsolute-like analysis

We were unable to run the DNA content analysis in scAbsolute using the original code base (see Supplement section S3 for details). Instead, we conducted an analysis of fragment overlaps similar in kind to the guidance offered by the vignettes in the scAbsolute repository (https://github.com/markowetzlab/scDNAseq-workflow/blob/main/vignettes/vignette-ploidy.pdf). In this section, we briefly summarize the approach used for comparison against scPlOver and ASCENT.

For each cell, we counted the fragment overlaps in each bin, then regressed these overlaps against the HMMCopy-assigned copy-number state *k* to obtain a predicted value for *k* = 2. We used a constrained quadratic regression in which the first two coefficients were restricted to be non-negative, and we required *f* (1) = 0 (i.e., the number of overlaps at state 1 should be 0) to avoid overfitting erroneous overlaps in *k* = 1 regions. We found this to perform better in practice than the `lmrob` robust linear regression used in scAbsolute and recommended in the vignette (likely due to the difference between actual fragment overlaps and the read density statistic used in the original publication; see Supplement section S3 for details). Note that despite the difference between these statistics, the inferred predicted values for copy 2 are strongly correlated with the published cell-level read density values (*R*^2^ = 0.948), especially for high-quality cells (Fig. **S1**A,B).

Then, at the individual level for each experimental dataset, we regressed the predicted overlaps at copy 2 against the average reads per copy for all G1 cells to obtain a reference curve. Finally, we computed the residual for each cell against this curve, and manually determined appropriate thresholds to classify a cell as not fitting well to this curve. Cells with excess positive residuals were called as higher DNA content than the initial CN assignment suggested, and cells with excess negative residuals were called as lower DNA content.

### 5.6 Simulating fragment overlaps

#### 5.6.1 Fixed-length

Given a bin size *L*, fragment length *𝓁*, number *n* of reads, and copy number *k*, we simulate sequencing fragments as follows. First, for each copy of the genome, we sample a random integer start position, and then tile the genome into fragments of equal length offset by the start value. This produces slightly less than *kL/𝓁* possible fragments. Then, we sample *n* fragments uniformly at random from the possible fragments. Finally, we iterate through pairs of sampled fragments, and count both the number of overlaps and the number of reference bases in each overlap.

#### 5.6.2 Variable-length

To incorporate realistic fragment length distributions rather than fixed fragment lengths, we sampled fragment lengths from the empirical fragment length distribution from an arbitrary cell (cell `128740A-R20-C25` from patient SPECTRUM-OV-037^7^). We constructed the empirical distribution by selecting those fragments with lengths between 50 and 1000 with a mapping quality of at least 20, then sampled fragment lengths proportional to their frequency. To divide a simulated bin into tiles, we first drew a large sequence of fragment lengths from the distribution (approximately 1000 times more than we would expect necessary to cover all copies of the bin). Then, we placed each fragment on a copy of the genome sequentially: first fragment 1 with length *𝓁*_1_ at start position 1 and end position *𝓁*_1_ + 1, then fragment 2 with length *𝓁*_2_ at start position *𝓁*_1_ + 2 and end position *𝓁*_1_ + *𝓁*_2_ + 2, etc. Once we reached a fragment that would overlap the end of the genome, this fragment was instead placed on the next copy of the genome. With the exception of the final fragment in each copy (whose effect is negligible when the average fragment length [here up to 1000] is much smaller than the bin size [here 500 000]), this approach produces a random tiling of the *k* genomic copies with fragment lengths distributed according to the empirical distribution. Then, we modeled observation of these fragments by sampling the desired number of fragments uniformly at random from these tiles. The number of overlaps and overlap bases were then quantified as in the fixed-length fragment case.

## 6 Declarations

### 6.1 Availability of data and materials

All sequencing data studied in this work was previously published and is publicly available in conjunction with the original publications: the FUCCI dataset [23] is available in the European Nucleotide Archive (ENA) under accession PRJEB61928; the dual marker datasets [27] are available in the National Center for Biotechnology Information (NCBI) Sequence Read Archive (SRA) under accession code PRJNA1158752; and the SPECTRUM HGSOC [9] processed data is available in Synapse (accession number: syn66366960) and raw data is available by requesting authorization to the Data Access Committee through dbGaP (accession number phs002857.v3.p1).

### 6.2 Code availability

scPlOver is available at https://github.com/shahcompbio/scplover. The Mondrian pipeline (https://github.com/mondrian-scwgs/mondrian_nf) was used to process data from FASTQ files, including alignment using `bwa-mem` [29] and CNA inference using HMMCopy [18].

### 6.5 Authors’ contributions

M.A.M. and G.S. developed the theoretical model. G.S. wrote the description of ASCENT and adapted its script for inclusion in benchmarking. M.A.M. developed the computational method, wrote the software, applied it to simulated and real data, and analyzed the results. M.A.M., G.S., and A.M. wrote the manuscript. A.M. and S.P.S. supervised.

## 6.6 Acknowledgments

We would like to thank the anonymous peer reviewers and the program committee of RECOMB-CCB 2026 for providing criticism which helped to improve this work.

## Footnotes

1 Neither BAF nor any other allele ratio statistic is informative for distinguishing *c*, as multiplying all allele-specific copy numbers by *c* does not change the ratios between the two parental alleles.

2 For example, a cell in G2 phase has the same ploidy as it had in G1 phase, but its DNA content *d* has doubled to 2*d*. A cell that undergoes WGD has both its ploidy and its DNA content *d* doubled to 2*d*.

3 We model overlap bases rather than overlaps as they provide a higher-resolution signal in general, which is especially useful in bins with few fragment overlaps.

4 While our model more naturally admits a discrete distribution like the Negative Binomial, unfortunately we are not aware of a closed-form solution for multivariate negative binomial regression with correlated variables [31, 32] which makes it expensive to optimize in the context of the Baum-Welch algorithm. We found it more valuable to model the strong covariance between reads and overlap bases than to model each dimension separately using counts.

5 Typically *R ≈* 2*F* in paired-end data, but differences in filtering for reads and fragments break this coupling.

6 This quality score combines sequencing and alignment metrics with goodness-of-fit measures from HMMCopy to produce a composite indicator of cell quality; see Laks et al. [25] for details.

7 Distributions appeared similar in shape for other cells from the same patient and for cells from the FUCCI [23] and dual marker [27] datasets.

