## Supplement for "scPlOver: inferring DNA content from amplification-free single-cell WGS using fragment overlaps"

### Supplemental materials for "scPlOver: inferring DNA content from amplication-free single-cell WGS using fragment overlaps"

#### S1 Derivation of total number $M$ of edges in overlap graph

Consider a region with length $L$ and $k$ copies, with fragment length $\ell$. Each of the $k$ copies of the region is divided into $L/\ell$ tiles, so there are a total of $T = kL/\ell$ possible tiles. Assuming that the read start sites are not exactly the same in any two copies, each tile overlaps two other tiles in every other copy – that is, each tile has $2(k-1)$ other tiles which, were the index tile and the other tile both observed, would represent an overlap. Because there are $T$ such index tiles, the number of possible overlaps is then $2T(k-1)\frac{1}{2}$ (where the latter $\frac{1}{2}$ avoids double-counting the same overlap). As a result, the total number of edges in the overlap graph is $M = 2T(k-1) * \frac{1}{2} = T(k-1)$.

#### S2 ASCENT formulation

In this section, we describe our understanding of the DNA content estimation module from the copy-number inference method ASCENT [1], which we extracted to test on data with experimentally controlled DNA content. After copy number calling, ASCENT evaluates whether a region's DNA content should be adjusted (doubled, halved, or kept the same) using a probabilistic overlap model.

**Model definitions:**

- $k$ — total observed number of overlaps.
- $c$ — sequencing coverage (per-base probability of observing a read).
- $d$ — expected duplication rate (noise term).
- $N$ — DNA content (number of chromosome copies).
- $f_N$ — mean fragment length for DNA content state $N$.
- $n_N$ — number of reads assigned to DNA content state $N$.
- $b_N$ — number of bins with $N$ copies
- $\ell$ — length of bin.

The model is based on per-position overlaps. In the absence of noise, given DNA content $N$, the probability of a given position on a distinct chromosome copy being sequenced is $c/N$. The number of distinct combinations of possible overlaps at a position is $\binom{N}{2}$. Thus, the expected number of overlaps at a position is

$$p_N = \frac{c}{N} \cdot \frac{c}{N} \cdot \binom{N}{2}.$$

The ASCENT model uses a duplication rate $d$ to account for noise (e.g., misalignment, incorrect copy number, index hopping), and computes an effective DNA content $(1+\frac{d}{2})N$, yielding an expected number of overlaps

$$p_N = \frac{c^2}{((1+\frac{d}{2})N)^2} \cdot \binom{(1+d)N}{2}.$$

Note that since $(1+d)N$ is non-integer the combinatorial term is computed using $\Gamma$ distributions as a continuous generalization of factorials. Also note that the probability term uses $(1+\frac{d}{2})$ while the combinatorial uses $(1+d)$ for reasons that are unclear to us, but this is consistent with the implementation in the ASCENT code base.

For a pair of fragments, the expected overlap size is computed as

$$\text{OS} = \frac{1}{2} \cdot \frac{\sum_N f_N \cdot n_N}{\sum_N n_N}.$$

which in the absence of systematic fragment length differences between DNA content states is approximately half of the mean fragment length. The expected number of overlaps over genomic span $b$ with bin length $\ell$ is

$$\lambda = \frac{\ell}{\text{OS}} \sum_N p_N \cdot b_N.$$

The likelihood of the observed number of overlaps $k$ is evaluated using a Poisson distribution:

$$P(k \mid \lambda) = \text{Poisson}(k \mid \lambda).$$

ASCENT computes the odds ratio of doubling or halving the DNA content relative to the predicted copy-number profile, and accepts the change if the odds ratio is less than 0.01.

**Additional Details** ASCENT downsamples to a fixed coverage of $0.005\times$ for all cells, such that the DNA content specific probabilities are constant for all cells.

**Filtering**

- Remove cells where $\geq 50\%$ of the genome lacks copy-number assignments.
- Remove reads whose start sites are within 1 bp of another read.
- Remove fragments below the 5th and above the 95th percentile of fragment length.
- Trim 5 bp from each end of every fragment.

### S3 scAbsolute

In this section, we describe our understanding of the scAbsolute [3] method derived from both the paper and the code base, and our challenges with running the original code.

#### S3.1 scAbsolute method

##### S3.1.1 Read density

The DNA content prediction module in scAbsolute uses a statistic called "read density" to quantify the extent of overlapping reads in each cell and bin. In the paper, this quantity is described as, alternatively, "the number of overlapping reads per region of the genome" (in Results), or "the average number of overlapping reads across genomic bins" (in Methods). The definition of read density is limited to these two plain English descriptions, and neither is described in sufficient detail to replicate. As such, reproducing this quantity requires examination of the code base.

First, duplicates are removed and fragments are counted from the BAM file (see `data/readPosition/extract-start-sites.sh`, specifically line 33). Then, the results are restricted to valid chromosomes and non-NaN values (see `data/readPosition/get_table.R`, specifically lines 45-53). Finally, this table is read into `R/scAbsolute.R`, but when it is passed to a `GRanges` object, the absolute value of the template length is used. This will incorrectly represent the rightmost read in each pair (with the larger position) as an additional fragment with the same length, effectively doubling the length of every fragment. Note that obviously this resulting value is strongly correlated with the value that would be derived from the correct coverage, and as such this bug is unlikely to account for a substantial change to the results of the original publication.

Given the table of fragment start positions and lengths, rather than counting overlaps or overlap bases, `R/scAbsolute.R` computes a modified coverage value for each bin by first using `GenomicRanges` to compute per-base coverage, then changing values of 1 to 0, and finally taking the base-level average using the binnedAverage function. The result is a sort of intermediate value between coverage and overlaps, as it does not count bases covered by only 1 fragment, but it scales linearly with the number of fragments covering a position rather than exponentially as the number of overlaps or overlap bases does (as the latter quantity counts each pair of fragments separately).

##### S3.1.2 Regression and classification

To incorporate fragment overlaps in DNA content estimation, scAbsolute computes two statistics for each cell:

- Predicted read density at copy-2 (confusingly, this value is also referred to as "read density"): the result of regressing the previously described "read density" statistic for all bins in each cell against the inferred copy-number state, and returning the value predicted by the regression for state 2.
- Average reads per copy $\rho$: for a given copy-number profile with fixed-width bins, $\rho = \frac{1}{N}\sum_{i=1}^{n}\frac{x_i}{c_i}$, where $x_i$ is the number of reads in bin $i$ and $c_i$ is the inferred copy-number state for bin $i$.

According to the original publication, computing these two statistics for cells with correctly-assigned DNA content results in a quadratic curve, which can then be used as a reference for other cells generated using the same sequencing platform. The authors propose selecting the DNA content value that minimizes the residual against this reference curve.

It appears that the read density statistic does not adequately account for fragment length. In the FUCCI cell-cycle-sorted data released with scAbsolute, fragment length is strongly related to DNA content (i.e., cell cycle; Fig. 3B). After accounting for fragment length, as demonstrated in principle on simulated data (Section 2.2, Fig. 2), the signal separating G2 cells from a quadratic curve based on G1 cells largely disappears (Fig. **S1**D,F) and the G1 and G2 cells are no longer clearly separated by the residual against the regression curve inferred from G1 cells (Fig. **S1**G).

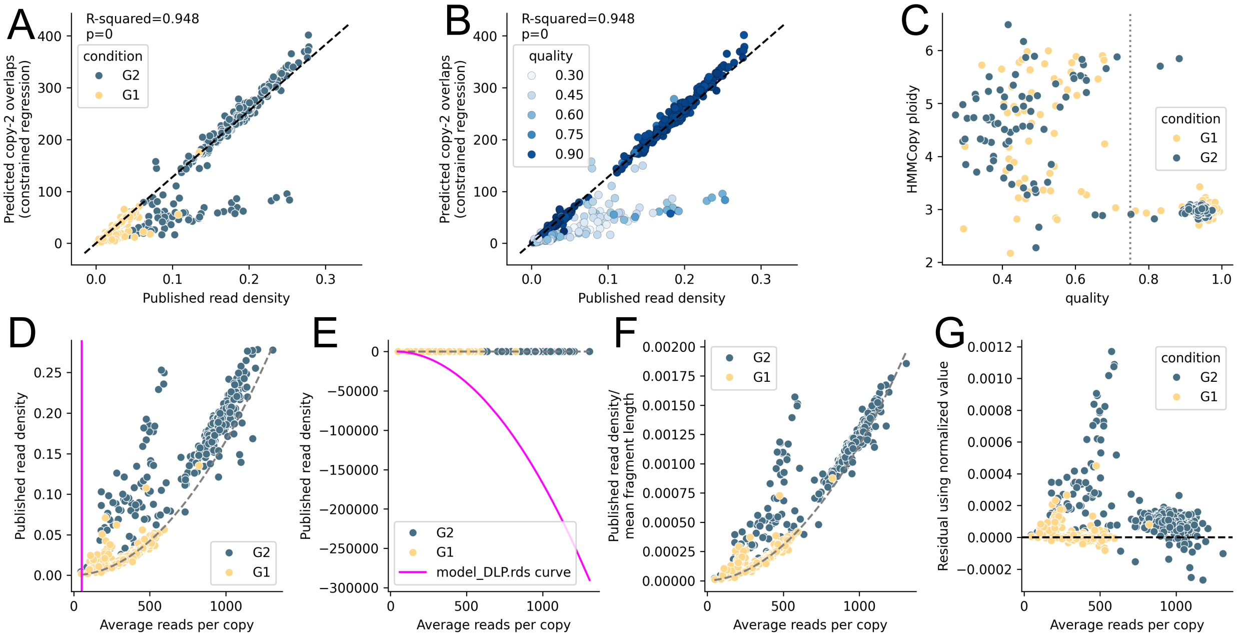


**Supplemental Fig. S1**: **scAbsolute [3] FUCCI dataset and published results. A.** Inferred copy-2 predicted overlaps from our implementation (y-axis) vs. published read density values from the original scAbsolute publication (x-axis), colored by cell cycle state. **B.** Same as A with cells colored by quality. **C.** Inferred HMMCopy ploidy (y-axis) vs. HMMCopy quality score (x-axis) for cells in the FUCCI dataset, colored by cell cycle state. Dotted gray line indicates quality cutoff of 0.75 which was used for supplemental analyses of FUCCI dataset. Cells are colored by cell cycle state. **D.** Published read density values (y-axis) as a function of the published reads per copy (x-axis) for experimentally sorted cells from the FUCCI dataset. Polynomial regression on G1 cells is shown in grey, and the polynomial curve included in the `model_DLP.rds` file is shown in magenta. Cells are colored by cell cycle state. **E.** Same values as B but with different axes to show the shape of the `model_DLP.rds` curve. **F.** Published read density values divided by mean fragment length (y-axis) as a function of the published reads per copy (x-axis) for experimentally sorted cells from the FUCCI dataset. Polynomial regression on G1 cells is shown in grey. Cells are colored by cell cycle state. **G.** Residuals of read density divided by mean fragment length against the curve (shown in F) inferred from G1 cells only (y-axis). Cells are colored by cell cycle state. Black dashed line indicates $y = 0$.

#### S3.2 Code base challenges

Unfortunately, while we were able to re-implement the cell-level read density regression and show that our implementation closely matches the results released alongside the original paper (Fig. **S1**A), we encountered a number of issues in trying to evaluate scAbsolute in practice.

1. In the original publication, the method for evaluating whether a cell's DNA content is assigned correctly is to compare cells to a reference curve generated via regression on a subset of DLP cells. However, at time of writing (May 1, 2026), the scAbsolute code base does not include the model file representing the curve to be used as reference for read density – the code to load this model (`https://github.com/markowetzlab/scAbsolute/blob/70ea6a8d3968a56d170bc7ca78db327c6fdcbcf1/R/scAbsolute.R#L624`) refers to a subdirectory `data/models` of the repository which does not exist. Exploring further, there is indeed a file `data/ploidy_prediction/model_DLP.rds`, but this curve does not appear to be in the same space as the published values despite matching variable names (Fig. **S1**D-E).
2. The recommended method for running scAbsolute is as part of the lab's "scDNAseq-workflow" pipeline, which is hard-coded to use the minimum error method for inferring DNA content instead of frag-

ment overlaps (see line `https://github.com/markowetzlab/scDNAseq-workflow/blob/7da985e793967c4c222773536d142760139ec29a/workflow/scripts/run_scAbsolute.R#L135`). If this line of code were changed, the pipeline would likely encounter an error as the model referred to in the code does not exist (see previous point).

##### S3.3 Reproducing scAbsolute-like analysis

While we were unable to compare against the scAbsolute method directly using the original code base, we used the guidance in a vignette in the repository to compare against the scAbsolute method in principle – we refer to this method as "scAbsolute-like". For each cell, we regressed read overlaps against assigned copy-number state, and returned the value predicted at copy-number state 2 to compare between cells. Then, for all cells in the dataset, we plotted these predicted copy-2 overlaps against the average number of reads per copy. Leveraging the experimental cell cycle labels, we obtained a reference curve by regressing copy-2 overlaps against average reads per copy in the G1 cells alone. Following the guidance in the vignette, we then inspected the residuals for each cell against this reference curve, identified an absolute threshold for distinguishing cells that were separated from the curve, and classified any cells exceeding this threshold as those whose DNA content should be doubled (with an excessive positive residual) or halved (with an excessive negative residual) with respect to the original ploidy value obtained by HMMCopy.

We tested a few different regression approaches to obtain predicted copy-2 overlaps. For this analysis, we restricted analysis to cells with an HMMCopy quality score of at least 0.75, as these showed the strongest correspondence between the published read density values and our regression (Fig. **S1**A-B) and had more consistent HMMCopy ploidy calls (Fig. **S1**C), which are fixed as input in this analysis. We found that a constrained quadratic regression (in which $f(1) = 0$ and the first two coefficients are non-negative) outperformed both robust linear regression ("lmrob" R package used in scAbsolute and the vignette) and the original published read density values in distinguishing G2 cells in the FUCCI dataset (accuracy 97.5% for constrained regression vs. 96.8% for lmrob and 93.4% for original read density; Fig. **S2** and Fig. **S3**A-D). We proceeded to apply this constrained quadratic regression approach to the two DM datasets [2] as well (Fig. **S3**E-L). In these datasets, there is little to no detectable separation between G2 and G1 phase cells, and as expected the residuals in this space perform poorly in identifying G2 cells (DM tetraploid, accuracy 41.7%; DM diploid, accuracy 45.7%). This is likely due to the fact that neither the published read density nor the predicted copy-2 overlaps used in this analysis controls for fragment length, and while the FUCCI dataset has a strong association between fragment length and cell cycle, this association appears weaker in the DM datasets (Fig. **S3**D,H,L).

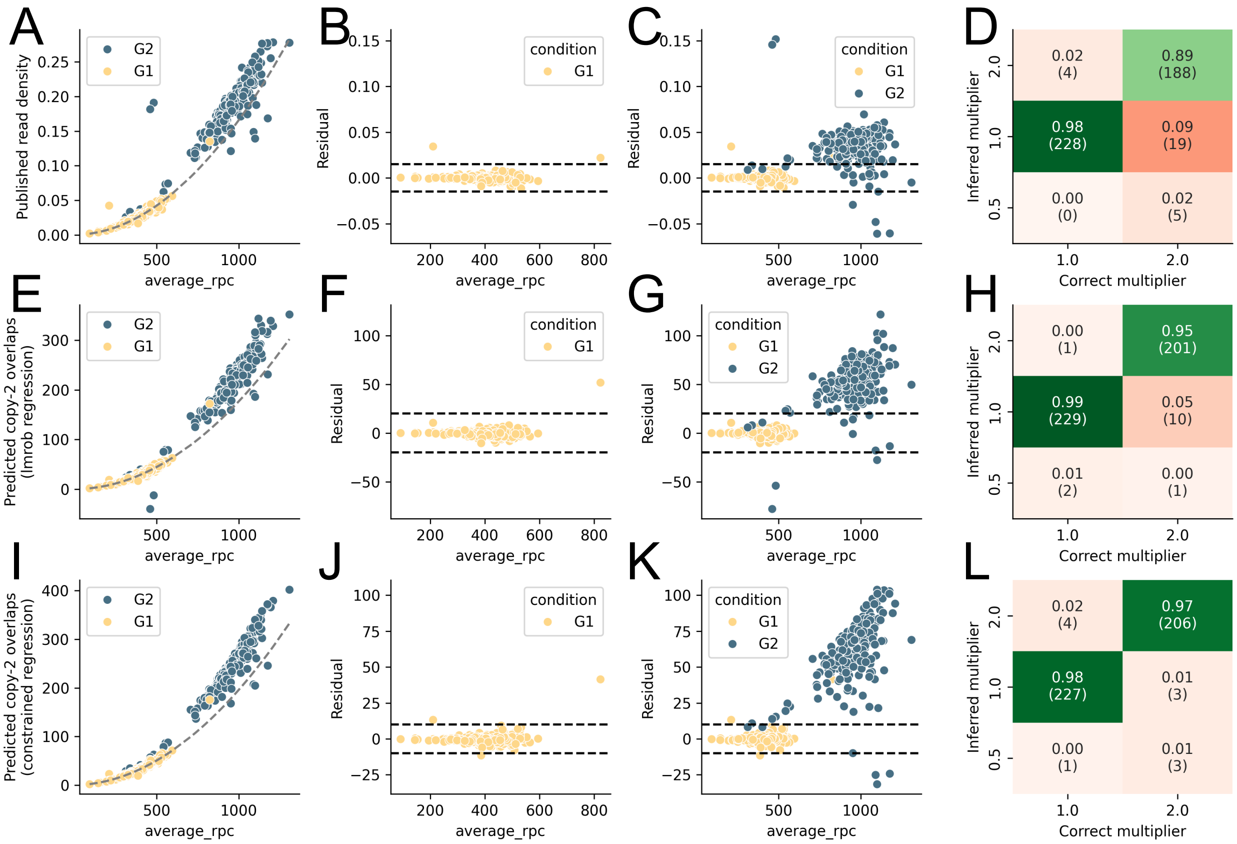


**Supplemental Fig. S2**: **Comparing cell-level regression approaches for copy-2 overlaps. A.** scAbsolute published read density values (y-axis) vs. average reads per copy (x-axis). Quadratic regression fit only to G1 cells is shown in gray (note that for A only, only those G1 cells with average reads per copy above 250 were included in the regression). **B.** Residuals of read density against the regression line shown in A. Dotted lines indicate manually identified cutoff to classify cells as doubled, halved, or appropriate DNA content. Showing G1 cells only. **C.** Same as B, but showing all cells. **D.** Confusion matrix comparing correct multiplier (x-axis) to inferred multiplier according to residuals (y-axis). Each entry is annotated with the proportion of cells in the column (i.e., in the "correct multiplier" category) as well as the raw cell count in parentheses, entries are colored according to correctness of the corresponding classification, and color intensity corresponds to value (colors scaled separately for all incorrect vs. all correct entries). **E-H.** Same as A-D but using the "lmrob" R package to predict the number of copy-2 overlaps in each cell (y-axis), as suggested in the scAbsolute vignette. Individual-level regression is fit to all G1 cells unlike in A. **I-L.** Same as A-D but using constrained quadratic regression ($f(1) = 0$, first 2 coefficients must be non-negative) to predict the number of copy-2 overlaps in each cell (y-axis). Individual-level regression is fit to all G1 cells unlike in A.

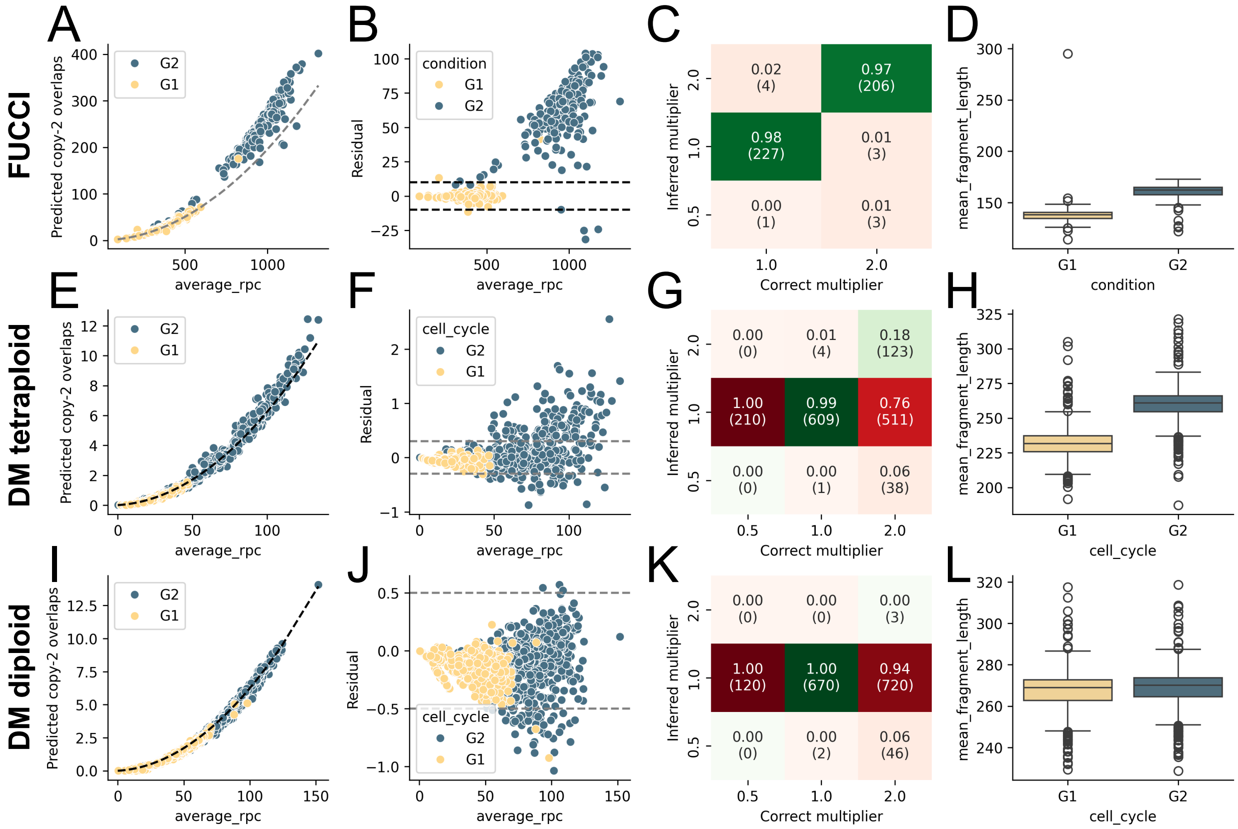


**Supplemental Fig. S3**: **Applying scAbsolute-like regression to experimental datasets.** **A.** Predicted overlaps at copy 2 from constrained quadratic regression (y-axis) vs. average reads per copy (x-axis) for cells in the FUCCI cell cycle sorted dataset [3]. Quadratic regression fit to G1 cells only is shown in gray. **B.** Residuals of read density against the regression line shown in A. Dotted lines indicate manually identified cutoff to classify cells as doubled, halved, or appropriate DNA content. **C.** Confusion matrix comparing correct multiplier (x-axis) to inferred multiplier according to residuals (y-axis). Each entry is annotated with the proportion of cells in the column (i.e., in the “correct multiplier” category) as well as the raw cell count in parentheses, entries are colored according to correctness of the corresponding classification, and color intensity corresponds to value (colors scaled separately for all incorrect vs. all correct entries). **D.** Mean fragment length (y-axis) vs. cell cycle state (x-axis) for cells in the FUCCI dataset. **E-H** Same as A-D for the double marker (DM) tetraploid dataset [2]. **I-L** Same as A-D for the double marker (DM) diploid dataset [2].

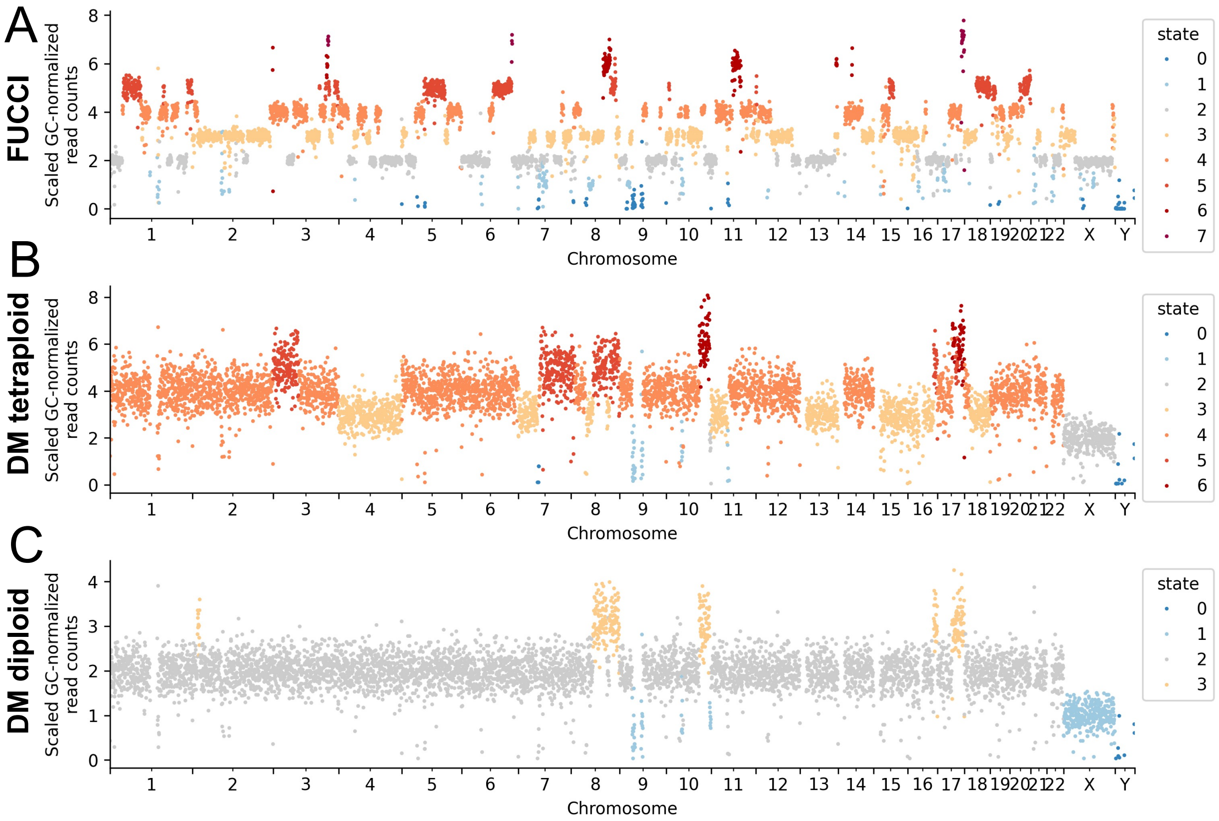


**Supplemental Fig. S4**: **Example G1 copy-number profiles for cells from experimental datasets. A.** Copy-number profile for an example G1 cell from the FUCCI dataset with quality $0.948$, ploidy $2.97$, and coverage depth $0.213\times$. Each point is a 500-kilobase bin colored by the assigned integer copy-number state from HMMCopy. The x-axis shows genomic position, and the y-axis shows the GC-normalized scaled read count for the bin. **B.** Copy-number profile for an example G1 cell from the DM tetraploid dataset with quality $0.946$, ploidy $3.68$, and coverage depth $0.030\times$. Plot details are the same as A. **C.** Copy-number profile for an example G1 cell from the DM diploid dataset with quality $0.976$, ploidy $1.95$, and coverage depth $0.027\times$. Plot details are the same as A.

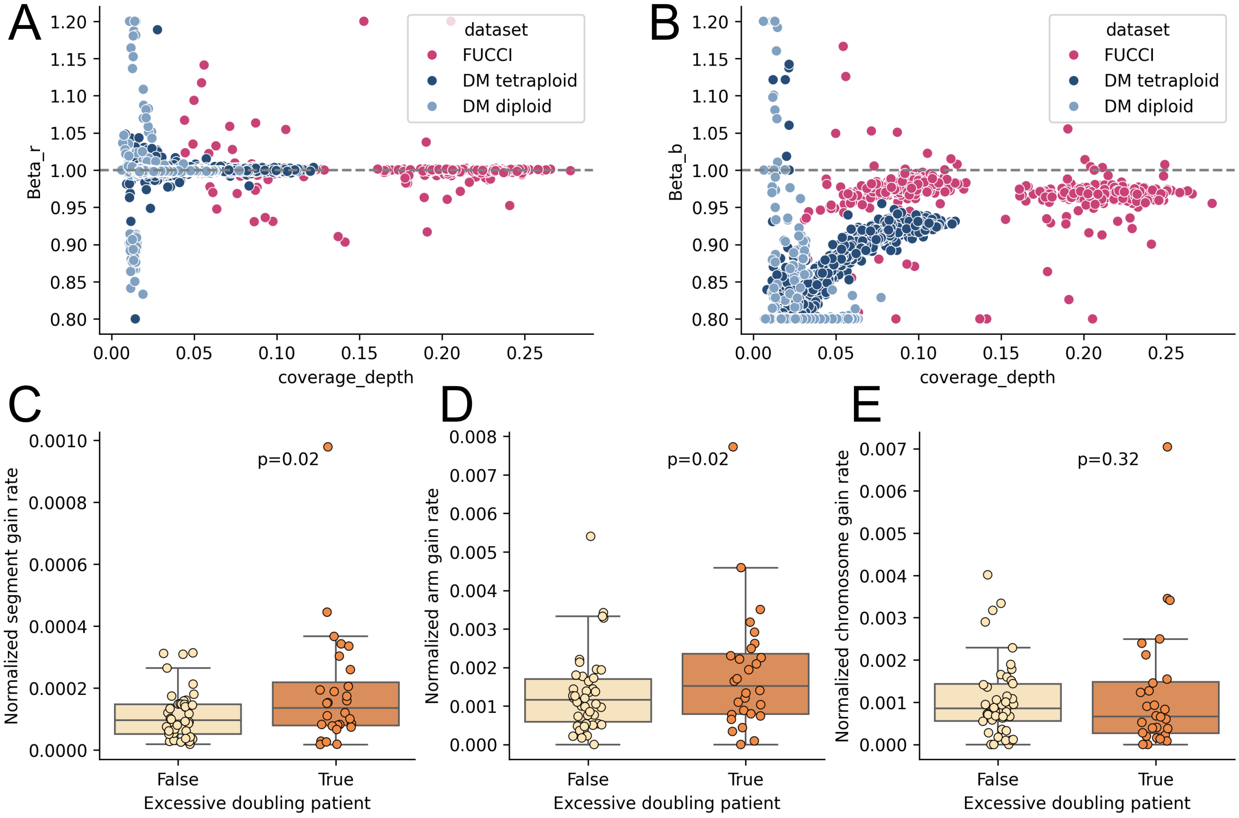


**Supplemental Fig. S5**: **Learned parameters $\beta$ for experimental datasets. A.** Inferred scaling factor $\beta_r$ for mean reads in the Gaussian emission model (y-axis) as a function of coverage (x-axis). Points are colored by dataset. Gray line indicates $y = 1$ (i.e., no scaling). **B.** Same as A but showing scaling factor $\beta_b$ for bases on the y-axis. **C.** Normalized rate of segment copy-number gains (y-axis) grouped by "excessive doubling" status (i.e., whether patient's proportion of doubled cells (called by scPlOver) exceeded the proportion of G2M cells in matched scRNA-seq) (x-axis). Each point is a sample, p-value for two-sided Mann-Whitney U test. **D.** Same as C for arm-level copy-number gains (y-axis). **E.** Same as C for chromosome-level copy-number gains (y-axis).
